# Adaptive Magnetic Resonance

**DOI:** 10.1101/2022.03.16.484410

**Authors:** Inbal Beracha, Amir Seginer, Assaf Tal

## Abstract

Nuclear magnetic resonance is one of the cornerstones of modern medicine and biomedical research. Over the past several decades, the speed and precision of in-vivo magnetic resonance imaging (MRI) and spectroscopy (MRS) have increased by leaps and bounds, by utilizing sophisticated excitation and acquisition techniques, from parallel imaging and compressed sensing to magnetic resonance fingerprinting. However, these approaches have all been static in nature, fixing measurement parameters in advance, in anticipation of a wide range of expected tissue parameter values, and are therefore sub-optimal for any given subject. We depart from the conventional framework of magnetic resonance and propose a new approach - termed adaptive magnetic resonance - which binds acquisition and excitation, by using the measured signal to update and fine-tune the measurement parameters in real time. This targets the specific tissue characteristics of the subject while they are being scanned. Adaptive magnetic resonance provides a completely new and previously-untapped avenue for improving the sensitivity and specificity of in-vivo magnetic resonance across all tissue contrast mechanisms. Equivalently, it can accelerate data acquisition compared to non-adaptive schemes, by obtaining the same precision using fewer, optimally tuned excitations. We demonstrate that an adaptive pulse sequence for measuring the transverse relaxation time (*T*_2_) of metabolites in-vivo improves upon the precision of static approaches by a factor of ≈ 1.7 - or, alternatively, accelerates acquisition 2.5-fold.

## Introduction

Since its inception, Magnetic Resonance has become ubiquitous in a wide range of disciplines, and has transformed many facets of biomedical research and medicine (1–4). In medical imaging, Magnetic Resonance Spectroscopy (MRS) and Imaging (MRI) provide objective measures of the composition and physiology of biological tissue, such as proton density (PD), longitudinal and transverse spin relaxation times (*T*_1_, *T*_2_), diffusion and magnetization transfer ratio, often in a quantitative manner. These metrics are powerful and platform-independent markers for a wide range of pathologies. At the heart of MRI and MRS lies the concept of a pulse sequence: a collection of radio-frequency and gradient pulses, which allows switching between contrast types by altering pulse shapes, amplitudes and delays. These sequence parameters - such as the echo time (TE), flip angle (FA), repetition time (TR) or inversion time (TI) - are static: They are set in advance, in anticipation of a particular type of contrast and a range of biological tissue parameters (e.g. *T*_1_ or *T*_2_), but are not varied between subjects.

We depart from the conventional static framework of MR and propose an adaptive approach to magnetic resonance, where incoming data is used in real time to update the sequence parameters. This approach operates by maintaining a probability distribution which quantifies our a-priori belief about obtaining different values for the tissue parameters of interest (e.g. *T*_1_). The distribution, *p*(*T*_1_,…) which starts out as uniform, gets updated each excitation, using the incoming signal to estimate the likelihood of different values of the tissue parameter. The updated distribution is used in turn to inform and sculpt the next excitation. This continuous, real-time interplay between the sequence and the acquired data, tailors the pulse sequence to the tissue parameters of the specific subject within the scanner. The update is carried out via a calculation which is both straightforward and computationally-efficient, allowing our adaptive approach to be applied to a wide range of contrast types. Our approach is also completely orthogonal to existing techniques, such as magnetic resonance fingerprinting (5–11), parallel imaging (12–15), compressed sensing (16–19) and model-based reconstruction (20, 21), and can be combined with each without impairing their performance.

The adaptive scheme is applied to the problem of measuring the transverse relaxation time *T*_2_ of n-acetyl-aspartate (NAA), a small metabolite found primarily inside neurons in the central nervous system. NAA is a powerful neuron-specific biomarker (22), and its transverse and longitudinal relaxation times change in response to changes in the neuronal microenvironment. NAA’s *T*_2_ has been demonstrated to vary in a wide range of neuropathologies, from multiple sclerosis to brain tumors and Alzheimer’s disease, and is often not measured in practice because of the low signal to noise ratio it exhibits (23). Using computer simulations, phantom measurements and in-vivo data, we show our adaptive approach yields approximately 1.5-2.0-fold improvement in sensitivity, without impairing its accuracy - or, equivalently, a two to four-fold reduction in experimental time. Specifically, we demonstrate that we can obtain the same precision for *T*_2_ in-vivo with only 40% of the data, accelerating acquisition 2.5-fold. Thus, our adaptive approach enables the fast and efficient quantification of the *T*_2_ time of NAA and its incorporation in clinical monitoring and prognostication.

## Methods

### Designing Adaptive Sequences

#### *T*_2_ *Estimation*

The well-established gold-standard for measuring *T*_2_ is a multiecho (MTE) experiment, in which the signal depends on the tissue (*T*_2_, PD) and sequence (TE) parameters as follows:

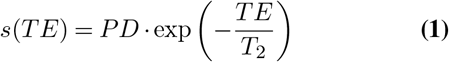

At the core of our adaptive approach is a probability distribution *p*(*T*_2_, *PD*) which quantifies the assumed a-priori probability of each value of the tissue parameters. Given such a distribution, we can estimate *T_2_* via 〈*T*_2_〉 = *∫ T*_2_*p*(*T*_2_, *PD*)*dT*_2_*dPD*, which can be used - alongside all previously selected TEs - to select the *N^th^* echo time *TE_N_* according to some predefined rule, *TE_N_* = *R*(〈*T*_2_〉, *TE*_1_,…, *TE*_*N*–1_). For example, as described below, we can set *R* to select the *TE* which minimizes the Cramer Rao Lower Bound (CRLB) for the estimated 〈*T*_2_〉. A new data point *s_N_* = *s*(*TE_N_*) is then acquired, and used to calculate the likelihood of each possible *T*_2_ and PD value given the new measurement: *L*(*T*_2_, *PD*|*s_N_*). If the data has Gaussian white noise with zero mean and variance *σ*^2^, then:

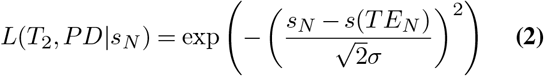

The standard deviation of the noise, *σ*, can be easily and accurately estimated from an initial fast excitation with all radiofrequency pulses turned off. The distribution *p*(*T*_2_, *PD*) is then updated via Bayesian estimation by multiplying the likelihood by the initial prior and normalizing, and the adaptive cycle can begin anew. This approach is illustrated in Fig. 1 for the simple one-dimensional case of a single tissue parameter, *T*_2_, keeping PD=1.

**Fig.1.**
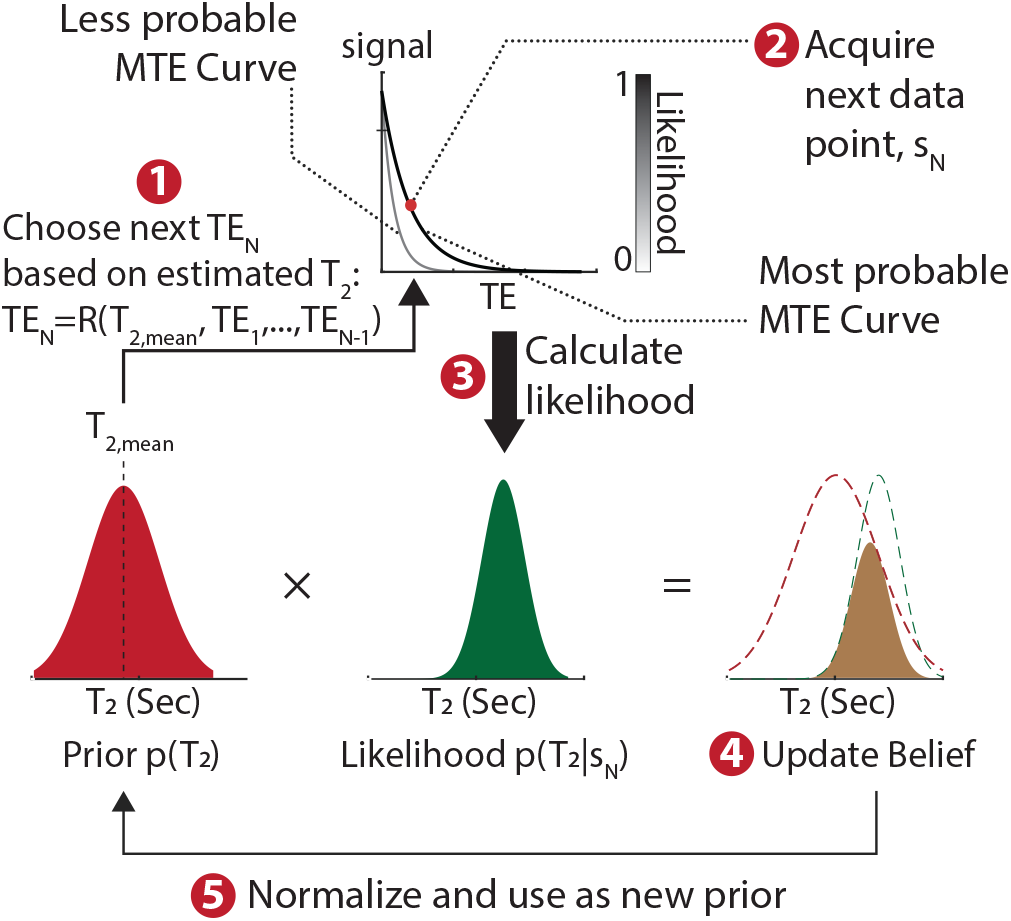
The adaptive measurement scheme starts out from a prior distribution *p*(*T*_2_) for the estimated parameter, *T*_2_, indicating how likely believe we each value of *T*_2_ is (initially *p*(*T*_2_) is uniform; we omit *PD* for simplicity). We choose our next TE using some predetermined rule *TE_N_* = *R*(*T*_2_, *TE*_1_,…,*TE*_*N*–1_), based on the current assumed value of *T*_2_, 〈*T*_2_〉 = *∫p*(*T*_2_)*T*_2_*dT*_2_, and previously measured echo times. New data is acquired at the chosen echo time, *s_N_* = *s*(*TE_N_*), and used to calculate the likelihood *L*(*T*_2_|*s_N_*) of each possible *T*_2_ value given the measured signal *s_N_*. Our belief is updated by multiplying the likelihood by the initial prior and normalizing; and this is used as the initial prior *p*(*T*_2_) for the next measurement.

#### Choosing the Update Rule

One possible real-time update rule *TE_N_* = *R*(〈*T*_2_〉, *TE*_1_…, *TE*_*N*–1_) chooses *TE_N_* in a way which approximates the optimal distribution of TEs for the estimated 〈*T*_2_〉 in the Cramer Rao Lower Bound (CRLB) sense. The CRLB provides a lower bound attainable by an unbiased estimator of *T*_2_, which is often attained in practice when using least squares fitting. Its minimization assures an optimal measurement, in the sense that it produces the highest possible precision (lowest standard deviation) for *T*_2_. For the MTE model in Eq. 1, the CRLB is a 2×2 covariance matrix for the parameter of interest *T*_2_ and the nuisance parameter PD. It is obtained by inverting the Fisher Information matrix, the elements of which are:

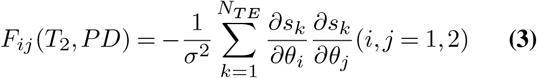

where *θ*_1_ ≡ *T*_2_, *θ*_2_ ≡ *PD*, *s_k_* ≡ *s*(*TE_k_*) and *N_TE_* is thenumber echo times acquired, equal to the number of excitations. One can calculate in advance the optimal distribution of echo times for each value of *T*_2_ and a fixed value of *N_TE_* by minimizing the second diagonal element of the CRLB matrix, which corresponds to the variance of *T*_2_. The rule *R* then chooses the next adaptive *TE_N_* such that the total distance (in, e.g., the *l*_1_-sense) of the distribution of measured and optimal echo times is minimal.

#### Inversion Recovery (IR)

As a second example, an IR acquisition obeys the signal model:

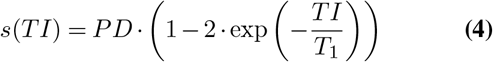

In IR, *T*_1_ is estimated by varying *TI*. An adaptive approach would use incoming data to update the 2D prior *p*(*T*_1_, *PD*), and choose each *TI* with an update rule designed in much the same way as for MTE (but using the CRLB matrix for the IR model in Eq. 4). A delay Δ is inserted at the end of each excitation to enable spins to return to thermal equilibrium; As the estimation of *T*_1_ improves, Δ can be shortened, by setting Δ = 5 · (〈*T*_1_〉 +3 · SD (*T*_1_)). Here, 〈*T*_1_〉 is the estimated mean value of *T*_1_, and SD (*T*_1_) is the standard deviation, both estimated via the 2D prior distribution. Three standard deviations are added as a safety margin, in case our estimation of *T*_1_ is inaccurate.

### Comparison to Static Approach

For *T*_2_ estimation, we have chosen to compare the precision (i.e. standard deviation) and bias (estimated minus true) of *T*_2_ for a fixed total acquisition time of two minutes. The static, non-adaptive approach must begin from the same prior belief, which in our case is a uniformly distributed between *T*_2_ ∈ [0.05, 0.45] seconds, discretized in *M* equispaced steps (*T*_2,1_,*T*_2,2_,…,*T*_2,*M*_). We calculated the average CRLB matrix over all values of *T*_2_ and minimized its second diagonal element - corresponding to the mean variance of *T*_2_ - to obtain a set of optimal echo times for the static case:

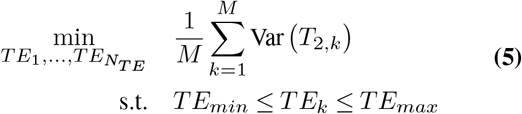

The optimization was carried out for *N_TE_* = 20. *TR* was fixed at 6 seconds for both the static and adaptive approaches, to allow the spins sufficient time to return to equilibrium, resulting in a total acquisition time *TA* of 120 seconds. Additional parameters were *TE_min_* = 0.04 sec, *TE_max_* =0.8 sec. This yielded a trimodal distribution of *TEs* which was sharply peaked at around 0.04, 0.11 and 0.37 seconds, at a ratio of 1:8:11.

For a static IR sequence, a similar optimization problem can be written down:

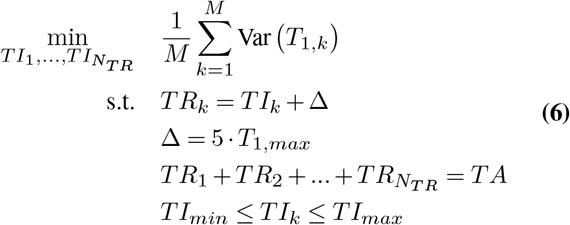

where Δ is a delay allowing the spins to return to thermal equilibrium (which also includes acquisition and water suppression). The optimization was carried out for all possible values of *N_TR_*, and the value of *N_TR_* which yielded the lowest mean variance was selected. The results of this optimization are shown in Fig. 2a (left) For *TI_min_* = 0.3 sec, *TI_max_* = 10 sec, *TA* = 120 seconds, *T*_1,*max*_ = 1.6 sec, *T*_1_ ∈ [0.6,1.6] sec.

**Fig.2.**
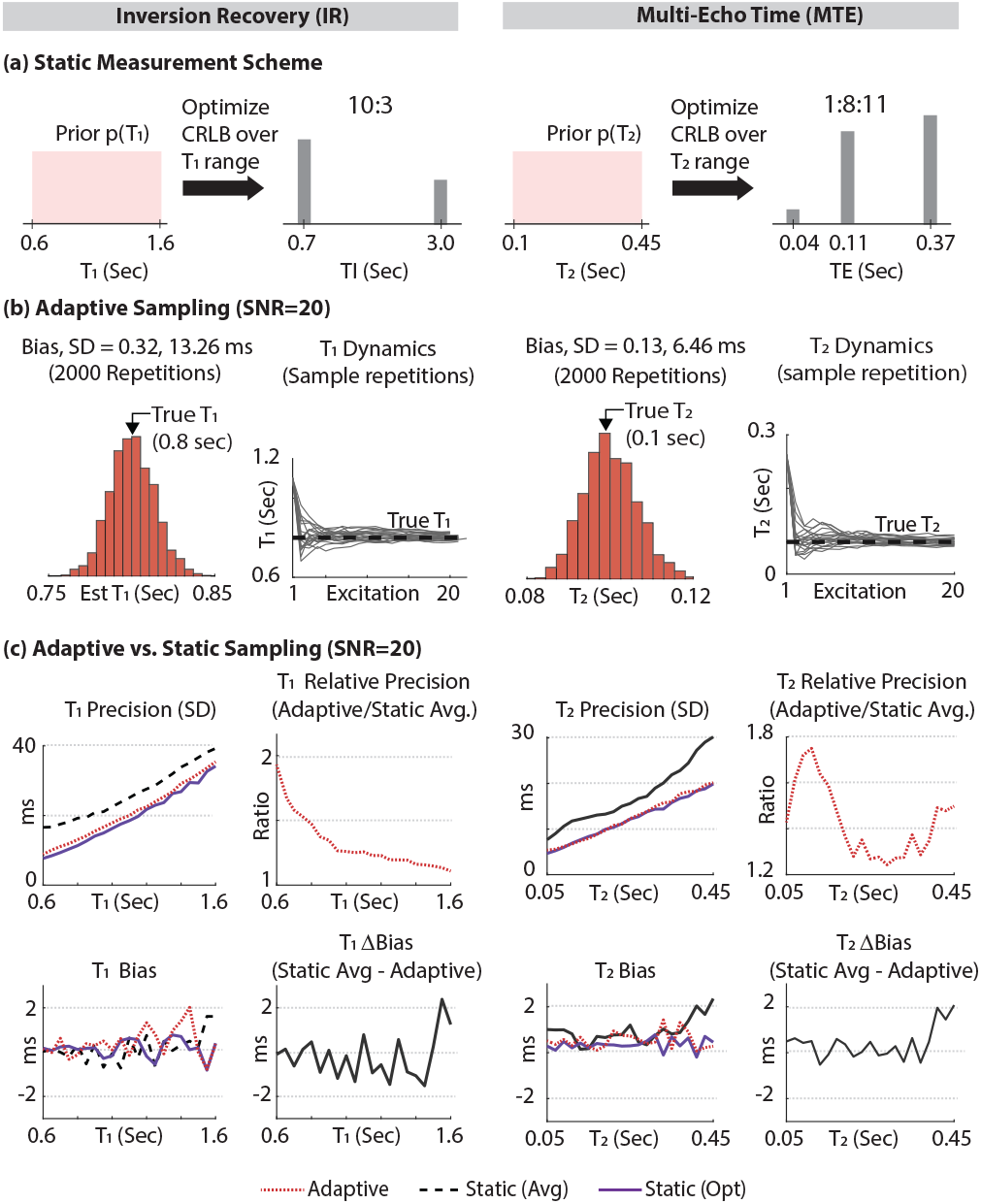
Simulation results for two widely used models: On the left, inversion recovery (IR) (Eq. 4), and on the right, a multi-echo (MTE) acquisition (Eq. 1). (a) The optimal TIs (TEs) which optimize the mean coefficient of variation of *T*_1_ (*T*_2_) over a range of prior values, *p*(*T*_1_) (*p*(*T*_2_)). These represent an optimal 120 second static acquisition. (b) The performance of the adaptive scheme for a specific *T*_1_=0.8 sec (*T*_2_=0.1 sec) for 2000 repeated iterations, each with different Gaussian noise, assuming SNR=20 for a single excitation at *TI*=∞ (TE=0 sec). Shown is the histogram of estimated *T*_1_ (*T*_2_) values from the 2000 iterations; its width is the standard deviation of *T*_1_ (*T*_2_), while the difference between its center and the true *T*_1_ (*T*_2_) is the bias. These histograms show the adaptive estimation is unbiased. Also shown are the temporal dynamics of *T*_1_ (*T*_2_) for a subset of 20 sample runs, showing that *T*_1_ (*T*_2_) converges to around its true value (dashed black line) very early on in the scan. (c) Comparison of the precision and bias of three experiments, using Monte Carlo simulations: The dashed black line corresponds to the optimal static experiment, with TIs (TEs) chosen as in (a). The dotted red line corresponds to the adaptive scheme. Finally, the solid purple line represents a static acquisition, with TIs (TEs) optimized for each specific single value of *T*_1_ (*T*_2_); this case is obviously unrealistic, since one does not know in advance the value of the parameter being estimated, but provides the best possible precision and bias one could hope to attain. These plots show the performance of the adaptive scheme (dotted red) is almost identical to that of the optimal (solid purple) approach. This is a result of the adaptive scheme converging very early on close to the true parameter values (as shown in (b)), resulting in near-optimal selection of TIs (TEs).

### Data Acquisition and Processing

All data was acquired on a Siemens 3T Prisma scanner (Siemens-Healthineers, Erlangen, Germany), using the manufacturer-provided 20 channel head coil for reception and the body coil for transmission, capable of maximal radiofrequency amplitude of 19 microtesla.

#### Sequence

We’ve used an in-house implementation of a Point-RESolved-Spectroscopy localization sequence. The adaptive protocol was implemented by using a real-time feedback mechanism built into the image reconstruction framework of Siemens (ICE), conventionally used for real-time navigators and on-the-fly image reconstruction. Real-time TEs and T2 estimations were stored in the log files for offline processing. The bandwidths of the excitation and refocusing pulses were 3.6 and 1.1 kHz, respectively, and their maximal amplitudes were both set at *γB*_1_ = 0.8 kHz. Water suppression took 426 ms and was achieved by applying a set of interleaved water-selective pulses and crusher gradients prior to excitation, with variable flip angles and delays optimized in-house.

#### Processing and Fitting

Even though the adaptive scheme employs real-time Bayesian statistics to inform its choice of parameters, once acquired its data can be processed using an identical pipeline to that of the static acquisition. This has been done for our simulations, and phantom and in-vivo datasets, to avoid differences relating to the processing pipeline. All processing used MATLAB 2021b (The Mathworks, Natick, MA). A singular value decomposition was used to combine the coils. Datasets were zero-filled four-fold along the spectral domain and apodized with a Gaussian window with a full-width at half maximum of 5 Hz. Phasing was achieved by summing all averages and maximizing the area underneath the real part of the spectrum between 0.5 and 4.2 ppm. Signal amplitudes used the maximum of the real part of the peak. Fitting employed a simple least-squares routine as implemented by MATLAB’s fminsearch function.

#### Phantom

Precision and bias of static and adaptive *T*_2_ estimations were validated on a spherical phantom, containing 1.9 L water doped with Gd-DTPA [0.5 mM] and metabolite concentrations similar to in vivo human brain: 12.5 mM NAA, 2 mM Choline chloride, 10 mM Creatine anydrous (Cr), 7.5 mM myo-Inositol (mIn), 10 mM L-Glutamtic Acid (Glu), 5 mM Sodium Lactate (Lac) and 2 mM *γ*-aminobutyric acid (GABA). Sodium Azide (*NaN*_3_, 0.01%) was added to avoid microbial growth. pH was adjusted by means of sodium phosphate buffer to be 7.2. For the phantom, prolonged 3.5-hour “gold-standard” MTE measurements yielded *T*_2_ =660, 545and 347ms for the NAA, Cr and Cho singlets at 2.01, 3.03 and 3.19 ppm, respectively. Because the *T*_2_s of NAA and Cr were longer than the assumed prior maximum of 450 ms, the adaptive scheme targeted the Cho peak (as opposed to the in-vivo measurements which targeted the NAA peak; see below).

#### Volunteers

A total of seven healthy volunteers were scanned after providing informed consent, as approved by Wolfson Medical Center Helsinki committee. Each volunteer was initially positioned head first supine. Following an anatomical *T*_1_-weighted MPRAGE scan, a 1.5 × 1.5 × 1.5 cm^3^ voxel was placed in parietal white matter and shimmed using in-house routines. Twelve adaptive and twelve static acquisitions were interleaved, each lasting two minutes (see below for justification of number of measurements). Total scan time was under 60 minutes.

#### Parameter Optimization

To optimize the acquisition for both the static and adaptive cases, one must begin with a uniform prior distribution on the parameter of interest (see Fig. 2). The range of possible outcomes will greatly affect the absolute and relative performance of each approach: wider a-priori ranges will penalize static approaches, but hardly affect adaptive ones. Therefore, a fair comparison must rely on literature reported ranges of the quantity of interest at 3T. Wyss et. al. (24) report a *T*_2_ range for the NAA singlet at 2.01 ppm of approximately 100-400 ms in healthy volunteers. NAA *T*_2_s as high as almost 500 ms have also been reported in the white matter of young adolescents (25). This is in line with a recent review which surveyed metabolite relaxation times at 1.5T and 3.0T across a wide range of pathologies and regions of interest (ROIs) (23). We have therefore chosen our prior on *T*_2_ to span [50,450] ms, as also depicted in Fig. 2. The same literature review also supports a range of in-vivo NAA *T*_1_s between [0.6, 1.6] seconds (23).

#### Statistical Analyses

##### Phantom

The standard deviation of *T*_2_ was computed for the static and adaptive approaches. The ratio of the static to the adaptive calculation was formed and the 95% confidence interval for ratio was calculated. A twotailed F-test was used to compare the variances of the two methods against the null hypothesis that they are equal. *In-Vivo:* A linear mixed model was used to test for statistically significant differences between the two approaches, as well as estimate the intra-subject standard deviation for each approach, using the model *T*_2_ ~ 1 + *method* + (1|subj). The linear mixed model can be used to obtain the intra-subject standard deviations, as well as their 95% confidence intervals.

#### Sample Size Justification

Sample sizes only depend on the ratio of the standard deviations being compared. For a phantom, given *m* adaptive and *n* static measurements, the 95% confidence interval (CI) (*α* = 0.05) on the ratio of two standard deviations is 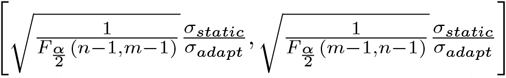, where *F* is the F-statistic. Setting *n* = *m* = 60 and assuming a ratio 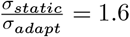 for Choline (based on simulations, using *T*_2_,*Cho*=347ms in the phantom), the CI is [1.24, 2.07]. For the in-vivo experiments we’ve used a Monte-Carlo simulation of our mixed-model approach, assuming 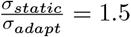, which is expected for the typical in-vivo white matter NAA *T*_2_s of around 350ms at 3T (24). This yielded a confidence interval of 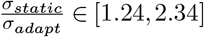.

## Results

### Simulations

Fig. 2 compares adaptive and static approaches for estimating *T*_1_ using IR, and *T*_2_ using MTE. Fig. 2a shows the optimal static TIs (TEs) which minimize the average coefficient of variation of *T*_1_ (*T*_2_) over a range of tissue parameter values, for a two minute acquisition. Fig. 2b shows the performance of the adaptive method for 2000 iterations of the adaptive approach, each taking 120 seconds and using different randomly generated Gaussian noise. Histograms show the distribution of the 2000 estimated *T*_1_s (*T*_2_s), indicating the approach is unbiased. The dynamics of *T*_1_ (*T*_2_) are also shown for a subset of the iterations, showing they converge very rapidly to the vicinity of the true value. Fig. 2c shows both approaches - static and adaptive - are unbiased, while the adaptive approach offers a 1.2-2.0 (1.3-1.8) fold increase in precision for IR (MTE), equivalent to a commensurate 1.4-4.0 (1.7-3.2) fold acceleration. Fig. 2 also shows the performance of the adaptive scheme (dotted red line) is almost identical to the best possible precision attainable, had we known in advance the true value of *T*_1_ (*T*_2_), and optimized the TIs (TEs) for that specific value in the CRLB sense. Such an optimal experiment is obviously unrealistic, since one does not know a-priori the true value of the estimated tissue parameter before running the experiment, but serves as a benchmark for the best possible outcome one might hope for.

Fig. 3 shows computer simulations which quantify how a bias in estimating *T*_2_ in real time (while the sequence is running) affects the precision and bias of *T*_2_, under the assumption the signal is correctly modeled without bias offline and refitted. Such bias might come about, for example, from sub-optimal estimations of the signal amplitude due to real-time time constraints. These simulations show that even a 20% (0.04 sec) real-time bias in *T*_2_ leads to a negligible ≤ 3% reduction in its precision and no discernible effect on its bias when correctly estimated off-line using the measured values.

**Fig.3.**
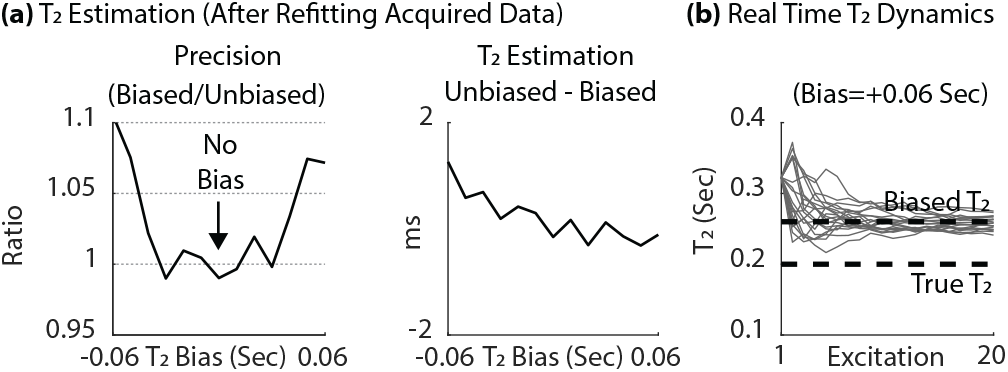
The effect of real-time *T*_2_ bias on its precision and bias when refitting offline, for when the true value of *T*_2_ is 0.2 sec. (a) Precision and bias of the *T*_2_ estimated off-line as a function of the bias in estimating *T*_2_ in real time, under the assumption that the signal is correctly modeled off-line using more advanced and computationally intensive model. Even large real-time biases of 20% result in almost no loss of precision or bias in the offline processing. (b) The real-time dynamics of *T*_2_ for a bias of 30% (+0.06 sec), showing the real-time acquisition converges to the biased value of *T*_2_ early on.

### Phantom

Fig. 4 shows experimental results from the phantom experiments, consisting of 60 repetitions of both the static and adaptive two-minute acquisitions. The adaptive acquisitions targeted the Choline singlet at 3.2 ppm (*T*_2,*Cho*_=347ms). Voxel placement is shown in Fig. 4a, and a sample single excitation is plotted in (Fig. 4b), showing typical spectral quality and SNR. The acquired data from a single static and adaptive acquisition is shown in Figs. 4c and 4d, with the fitted MTE curve (Eq. 1). Note how the static data points are acquired at the optimal TEs shown in Fig. 2a, while the adaptive data points are distributed unequally between the minimal TE (40ms) and ≈ 1.28 · *T*_2_; This distribution approximates the optimal one for specific value of *T*_2,*Cho*_=347ms of Cho, and was encoded into the update rule *R* of the adaptive sequence (the first step in Fig. 1). Both the adaptive and static acquisitions provide unbiased estimated of the true *T*_2_ (Fig. 4f), but the adaptive acquisitions yield better 1.76-fold better precision: *σ_adapt_* = 50 ms vs. *σ_static_* = 88 ms, with corresponding confidence intervals 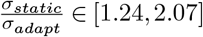. An F-test shows the ratio of these standard deviations to differ from unity in a statistically significant manner.

**Fig.4.**
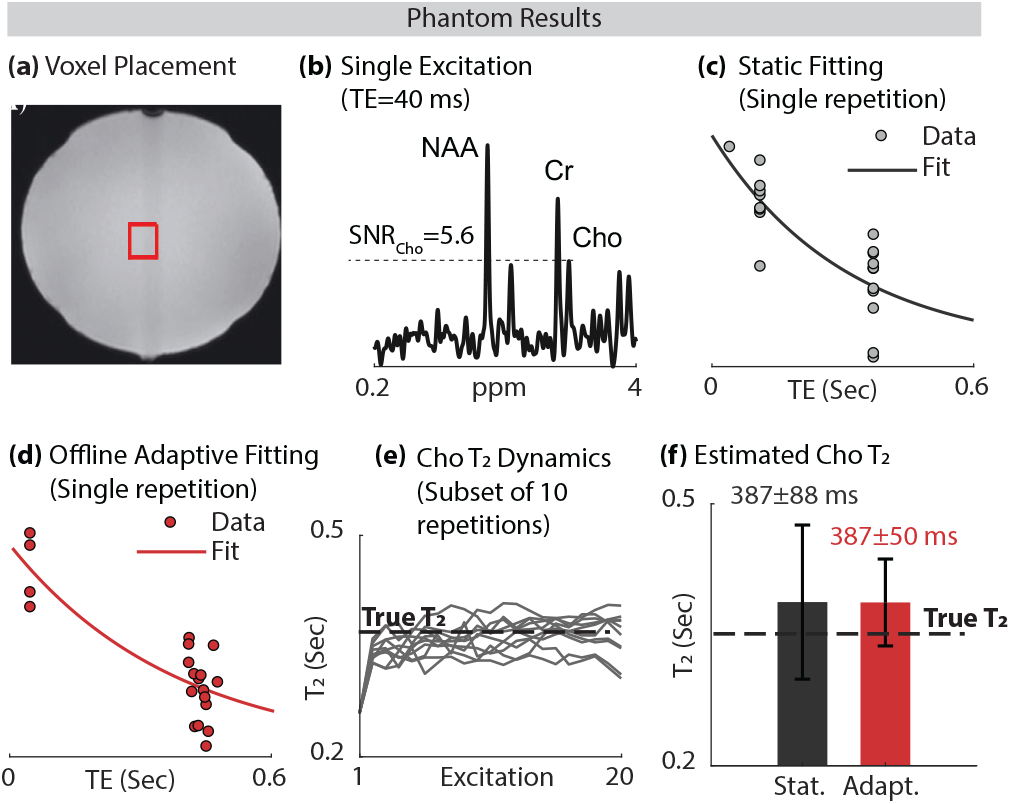
Phantom data. (a) 1.5 × 1.5 × 1.5 cm^3^ voxel placed at the center of the phantom. (b) Sample spectra: single excitation with water suppression, shown for a single excitation (TR/TE=6000/40ms). The SNR of the choline peak is 8.1. (c, d) Sample data and fitted MTE curve from a single repetition. Each acquisition consists of 20 data points, acquired over a two-minute interval. The static data has been sampled at the optimal echo times (shown in Fig. 2a) TE=40, 110, 370 ms, at a ratio of 1:8:11. The adaptive data has its TEs distributed around TE=40, 520 ms at an approximate 1:4 ratio, which is the optimal distribution of TEs (in a Cramer-Rao sense) corresponding to the true *T*_2,*Cho*_=347ms. (e) The dynamics of *T*_2_ for a subset of 10 repeitions out of the 60 in total. (f) Bars plots of *T*_2_ estimations (mean±SD over all 60 results). Both methods are unbiased, and the adaptive method exhibits better precision by a factor of 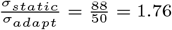 in line with the simulation predictions for *T*_2,*Cho*_=347ms.

### In-Vivo

Fig. 5a-e showcases sample data from a sample volunteer, including voxel placement, a typical single excitation at TE=40 ms, fitted MTE curves for a single repetition (out of the 12 repetitions acquired), and the dynamics of *T*_2_ estimation for all 12 repetitions. The adaptive acquisition targeted the NAA singlet at 2.01 ppm. Fig. 5f shows the distribution of fitted *T*_2_ values of NAA for each of the seven volunteers, providing visual indication the means are not significantly different in the statistical sense, and that the standard deviation of the adaptive scheme is overall better than that of the static one. The ratio 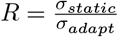 of standard deviations for each volunteer is shown above each bar plot. The mean ratio of standard deviations, averaged over all volunteers, is 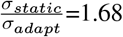, i.e. approximately 70% better for the adaptive approach. The mixed model analysis confirmed no statistically significant differences between the means of the two groups, and provided the following means and 95% CIs for the standard deviation of *T*_2_ for each approach: *σ_static_*=49.5 ms, *CI_static_* = [41.6, 59.0] ms, *σ_adapt_*=29.0 ms, *CI_adapt_* = [24.3,34.5] ms. The 95% CI for the ratio of the two standard deviations was [1.21,2.42], confirming the adaptive approach does indeed offer a statistically significant improvement in precision over the static one.

**Fig.5.**
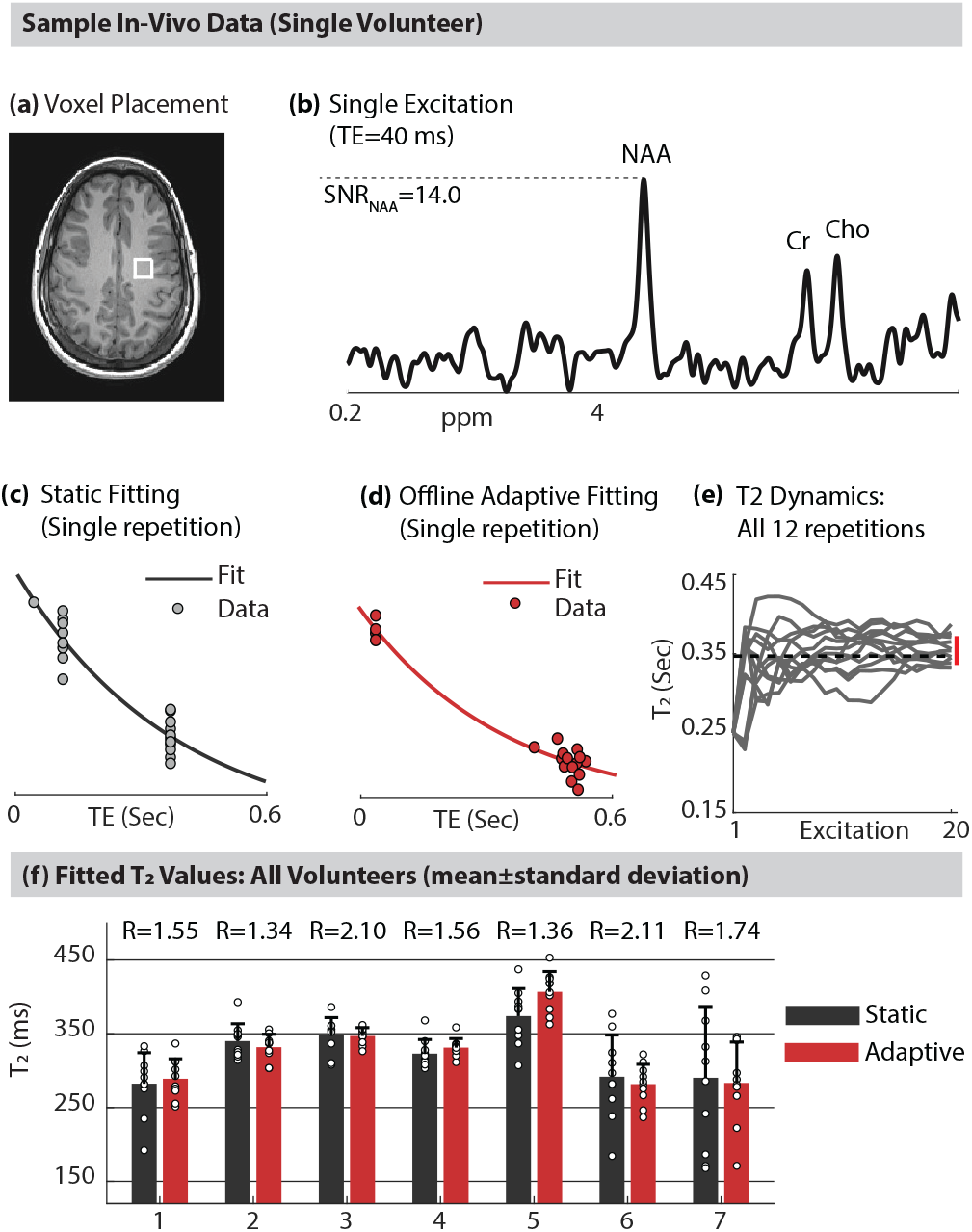
In vivo data. (a) Sample voxel placement (1.5 × 1.5 × 1.5 cm^3^) in parietal WM. (b) Sample single excitation (TR/TE=6000/40ms), showing *SNR_NAA_*=8.1. (c, d) Fitted MTE curves for two sample two-minute repetitions from the sample volunteertwo-minute acquisitions, showing how the echo time converges early on to the correct value. (e) Distribution of *T*_2_ estimations from both the static and adaptive approaches, showing the adaptive approach exhibits a statistically significant ≈ 1.9-fold improvement in its standard deviation. (f) Bar plots showing the mean and standard deviation of *T*_2_ for each method for each volunteer (circles represent estimations from each of the 12 repetitions). The number *R* above each pair of bar plots is the ratio of the static and adaptive standard deviations of *T*_2_ for each subject. Measurements are unbiased relative to each other, but the precision of the adaptive approach is approximately 70% better than the static one.

To determine the commensurate acceleration offered by the adaptive approach, we recalculated the mean precision *σ_adapt_* of the adaptive MR dataset by taking successively fewer excitations (omitting later excitations), until further removal of excitations resulted in worse precision by the adaptive approach compared to the static approach (*σ_adapt_* > *σ_static_*). The ratio of the number of retained excitations at this point, to the total number of excitations (20), was 8/20 = 0.4. Thus, our adaptive dataset confirms we could have attained the same precision by only using 40% of the excitations, corresponding to a 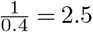-fold acceleration in measurement time.

## Discussion and Conclusions

Adaptive MR improves tissue parameter estimation in two ways. First, by informing the selection of optimal sequence parameters (TR, TE, TI, FA, etc) while the sequence is being executed, given our changing a-priori knowledge about the tissue parameters (*T*_1_, *T*_2_, *PD*, etc). Second, in some paradigms, updating our prior knowledge of the tissue parameters allows us to reduce total measurement time (or, equivalently, increase the number of measurements for a fixed total measurement time). For example, in IR, the post-excitation delay Δ needed for spins to return to thermal equilibrium. In the static case, this is often chosen to be five times as long as the longest projected *T*_1_ in the sample. However, an adaptive approach produces ever-increasing precise estimations of the true *T*_1_ in question; if the imaged tissue has a true *T*_1_ which is substantially shorter than the a-priori maximal assumed *T*_1_, Δ can be dramatically shortened in real time and, consequently, more averages can be acquired for a fixed total acquisition time. Our simulated, phantom and in-vivo data indicates that tissue parameter estimates converge very rapidly to their true values (Figs. 4e, 5e, 2b), which means that the sequence parameters also converge early on to near their optimal values.

The improved precision offered by adaptive schemes, along with the fast convergence of tissue parameters to their true values, means *N*-fold greater precision of adaptive sequences can be traded near-optimally for an *N*^2^-fold reduction in total acquisition time - up to a four-fold acceleration for IR and three-fold for MTE, given the suggested maximal 2.0-fold and 1.8-fold improvements to precision in simulations (Fig. 2c). Our adaptive in-vivo data produces the same precision as the static data with only 40% of the excitations, corresponding to a 2.5-fold acceleration. Because adaptive schemes maintain a distribution of the tissue parameters - e.g., *p*(*T*_2_, *PD*) for MTE - this can be used in real time to estimate the precision of the parameter of interest and use it as a stopping criterion, identifying cases in which convergence is poor and potentially prolonging the acquisition. Poor convergence can come about, e.g., due to patient motion or some other artifact which invalidates the assumed signal model.

The use of Bayesian estimation provides an efficient framework for incorporating prior knowledge into parameter estimation in real time. In the case of a signal with a simple Gaussian or other well-modeled noise distribution, it only involves the calculation of a likelihood function and the multiplication of two probability distributions. This might become somewhat time-consuming if the signal model contains multiple parameters, in which case these probability distributions become multidimensional and might require a substantial amount of memory and CPU time to store and multiply. To overcome this, they can be modeled using a low dimensional representation - e.g., sum of Gaussians - which can greatly enhance computational efficiency. Even using our brute-force 2D representation, we have timed the adaptive update stage to require approximately 35 ms. In cases where such calculations are too time-consuming to carry out within a single *TR*, the sequence can be partitioned into acquisition windows consisting of several *TR*s, and each window can be processed in bulk, allowing for more time to carry out computations. Alternatively, extremely fast neural-network solvers now exist which can extract tissue parameters from the acquired signal within milliseconds, making it possible to consider adaptive schemes even for complex problems such as magnetic resonance fingerprinting, or multi-pool magentization transfer models (20, 26–32). Neural networks can be used even in cases where a closed-form explicit model for the signal (e.g. Eq. 4 or Eq. 1) does not exist.

Any adaptive scheme must choose optimize some target function. In our in-vivo data, we updated the measurement parameter, *TE*, to minimize the variance in *T*_2_ for a single peak, NAA. In more complex scenarios one must carefully consider what to optimize. For example, in a BOLD-fMRI experiment, the echo time could be adjusted to maximize BOLD contrast between rest and activation based on each individual’s hemodynamic response. When imaging a tumor, contrast between the tumor margins and surrounding tissue on a *T*_1_-weighted image might be crucial, and therefore the contrast-to-noise ratio between two regions of interest in the image might be appropriate; These regions could be defined in advance, based on a quick scout image, or even identified in real-time using fast-performing neural networks during acquisition as part of the update process. Other optimization targets might modify the sampling in k-space based on whether the signal exhibits sufficient sparsity in some representation. The concept of adaptive sequences need not be confined to a single contrast type. When a pathology exhibits change in several contrast mechanisms, one can start out using a sequence capable of measuring more than one contrast type - such as a magnetic resonance fingerprinting acquisition with a highly variable schedule - and evaluate in real time which contrast type yields the greatest contrast-to-noise ratio, or deviation from some average healthy brain template. The sequence can then be adjusted not just to maximize contrast, but also to pinpoint and focus on the best type of contrast.

### Relation to Other Literature

The problem of adapting a process to incoming data in real time has been explored extensively within the field of signal processing; its best known implementation is the Kalman filter, which updates the state of a system based on its dynamical model and sensor data (33). Bayesian statistics provide an easy, straightforward framework for such updates. Indeed, adaptive Bayesian-based estimations have been used for measurements involving nitrogen vacancy centers, where they have been used to improve the precision of quantum sensors in estimating resonant spin frequencies in the presence of noise (34–37). The generality of the framework outlined in Fig. 1 makes it suitable for many sub-fields of NMR, from protein structure determination to solid state measurements. Adaptive schemes are also completely independent from other sources of acceleration - such as compressed sensing, magnetic resonance fingerprinting and parallel imaging - and can therefore be combined with them without detracting from their performance.

Several other real-time frameworks exist for MR imaging. Real-time estimation and prospective correction of subject motion is a major field of research in MR imaging, and has been approached using by acquiring navigator data within the sequence or using external field sensors (38–42). While both approaches share similarity in being real-time, prospective motion correction accounts for rigid body kinematic motion by updating the orientation of the gradients’ logical axes, but does not rely on the details of the pulse sequence used to acquire the image contrast, nor alter the sequence to change the contrast itself. Real-time MRI (43–45), another sub-field which rapidly acquires image slices at high (10-50 ms) temporal resolutions, contains no adaptive component to it at all.

## ACKNOWLEDGEMENTS

This work was supported by the Israeli Science Foundation Personal Grant 416/20. Assaf Tal acknowledges the support of the Monroy-Marks Career Development Fund and the historic generosity of the Harold Pearlman Family.

